# Complete and haplotype-specific sequence assembly of segmental duplication-mediated genome rearrangements using CRISPR-targeted ultra-long read sequencing (CTLR-Seq)

**DOI:** 10.1101/2020.10.23.349621

**Authors:** Bo Zhou, GiWon Shin, Stephanie U. Greer, Lisanne Vervoort, Yiling Huang, Reenal Pattni, Marcus Ho, Wing H. Wong, Joris R. Vermeesch, Hanlee P. Ji, Alexander E. Urban

**Author notes:** Co-corresponding authors **Corresponding authors:** Alexander E. Urban, Ph.D, Hanlee P. Ji, MD. These authors contributed equally (Bo Zhou, GiWon Shin).

## Abstract

We have developed a generally applicable method based on CRISPR/Cas9-targeted ultra-long read sequencing (CTLR-Seq) to completely and haplotype-specifically resolve, at base-pair resolution, large, complex, and highly repetitive genomic regions that had been previously impenetrable to next-generation sequencing analysis such as large segmental duplication (SegDup) regions and their associated genome rearrangements that stretch hundreds of kilobases. Our method combines *in vitro* Cas9-mediated cutting of the genome and pulse-field gel electrophoresis to haplotype-specifically isolate intact large (200-550 kb) target regions that encompass previously unresolvable genomic sequences. These target fragments are then sequenced (amplification-free) to produce ultra-long reads at up to 40x on-target coverage using Oxford nanopore technology, allowing for the complete assembly of the complex genomic regions of interest at single base-pair resolution. We applied CTLR-Seq to resolve the exact sequence of SegDup rearrangements that constitute the boundary regions of the 22q11.2 deletion CNV and of the 16p11.2 deletion and duplication CNVs. These CNVs are among the strongest known risk factors for schizophrenia and autism. We then perform *de novo* assembly to resolve, for the first time, at single base-pair resolution, the sequence rearrangements of the 22q11.2 and 16p11.2 CNVs, mapping out exactly the genes and non-coding regions that are affected by the CNV for different carriers.

## INTRODUCTION

Although “whole-genome” sequencing analysis is now routine practice, large fractions of the human genome that are complex, repetitive, or highly polymorphic, such as centromeres, telomeres, and segmental duplications, are still impenetrable to genome analysis. In even the most recent version of the human reference genome, these regions are left as “reference gaps” where no reliable reference sequence is available. There are a variety of reasons for this, one of which is that certain genomic regions are so structurally polymorphic across different individuals that it is impossible to collapse them into a single haploid reference (Demaerel et al., 2019). This is especially true for segmental duplication (SegDup) regions which make up approximately 5% of the human genome and comprise of hundreds of kilobases of highly polymorphic and repetitive sequences (Bailey et al., 2001, 2002). SegDups form the boundaries and also mediate the formation of recurrent chromosomal aberrations that are strongly associated with human diseases (Zhang et al., 2009; Stankiewicz and Lupski, 2010; McDonald-McGinn et al., 2015; Harel and Lupski, 2018). The high-degree of sequence homology between different SegDup regions precludes them from being resolved by short-read, linked-read, and even long-read sequencing (<30 kb), because of extensive ambiguous or inconclusive mapping of reads (Guo et al., 2016). Here, to overcome this technological challenge, we developed a generally applicable method CRISPR/Cas9-targeted ultra-long read sequencing (CTLR-Seq) which combines *in vitro* Cas9-mediated cutting of the genome and pulse-field gel electrophoresis, also known as CATCH (Jiang et al., 2015; Shin et al., 2017; Gabrieli et al., 2018), to isolate intact, haplotype-specific target fragments and to completely resolve, at base-pair resolution, large, complex, and highly repetitive genomic regions previously impenetrable to sequencing analysis.

Recurrent, SegDup-mediated heterozygous chromosomal aberrations in the form of large deletion and duplication copy number variants (CNVs) have the highest penetrance for neuropsychiatric disorders (Stankiewicz and Lupski, 2010; Sullivan et al., 2012). For example, the 22q11.2 Deletion Syndrome (22q11DS) is by far the highest known risk locus for developing schizophrenia, increasing the likelihood by at least 25-fold; 22q11DS patients also frequently develop autism, attention-deficit hyperactivity disorder, learning disabilities, heart diseases, immune system dysfunction, and hormonal deficiencies (McDonald-McGinn et al., 2015). Reciprocal deletions (MIM 611913) and duplications (MIM 614671) at 16p11.2 are strong risk factors for schizophrenia (SZ) (McCarthy et al., 2009) and autism spectrum disorder (ASD) (Sanders et al., 2012). Duplications produce a 14.5-fold increase in risk for SZ (Deshpande and Weiss, 2018; Miller et al., 2015), but ASD is frequent in carriers of deletions (22%) as well as duplications (26%) (Sebat et al., 2007; Marshall et al., 2008; Weiss et al., 2008; Walsh and Bracken, 2011; Niarchou et al., 2019). After the 22q11.2 deletion, the 16p11.2 deletion is the second most commonly identified microdeletion in clinical testing (Kaminsky et al., 2011). Similar to 22q11DS (Malhotra and Sebat, 2012; Kirov, 2015; Deshpande and Weiss, 2018), the 16p11.2 CNVs are also associated with broad developmental consequences including speech/language delay, intellectual disability, developmental delay, and ADHD; ~50% of either deletion or duplication carriers have at least one psychiatric diagnosis (Ghebranious et al., 2007; Green Snyder et al., 2016; Steinman et al., 2016; Bernier et al., 2017; Niarchou et al., 2019). Due to their strong associations with neuropsychiatric disorders, the 22q11.2 and 16p11.2 CNVs serve as key points of entry for the investigations of molecular etiologies of neurodevelopmental disorders (Blumenthal et al., 2014; Migliavacca et al., 2015; Deshpande et al., 2017; Ward et al., 2020).

The 16p11.2 CNVs are flanked by Break Point regions BP4 and BP5 (Zufferey et al., 2012) (Fig 1), which consist of >100 kb segmental duplications (SegDup) with >99% similarity to each other (Bailey et al., 2002). The duplicated sequences between BP4/5 are placed in direct (non-inverted) orientation which act as substrates for rearrangement creating either deletion or duplication (Bailey et al., 2002; Zufferey et al., 2012). Similarly, the 22q11.2 deletion CNV is also flanked by hundreds of kilobases of SegDup regions, also called low copy repeats (LCR), LCR22A and LCR22D (Figure 1A). These two large SegDup regions LCR22A and LCR22D rearrange and gives rise to the typical 3 Mb 22q11.2 deletion (Morrow et al., 1995; Edelmann et al., 1999a).

**Figure 1.**
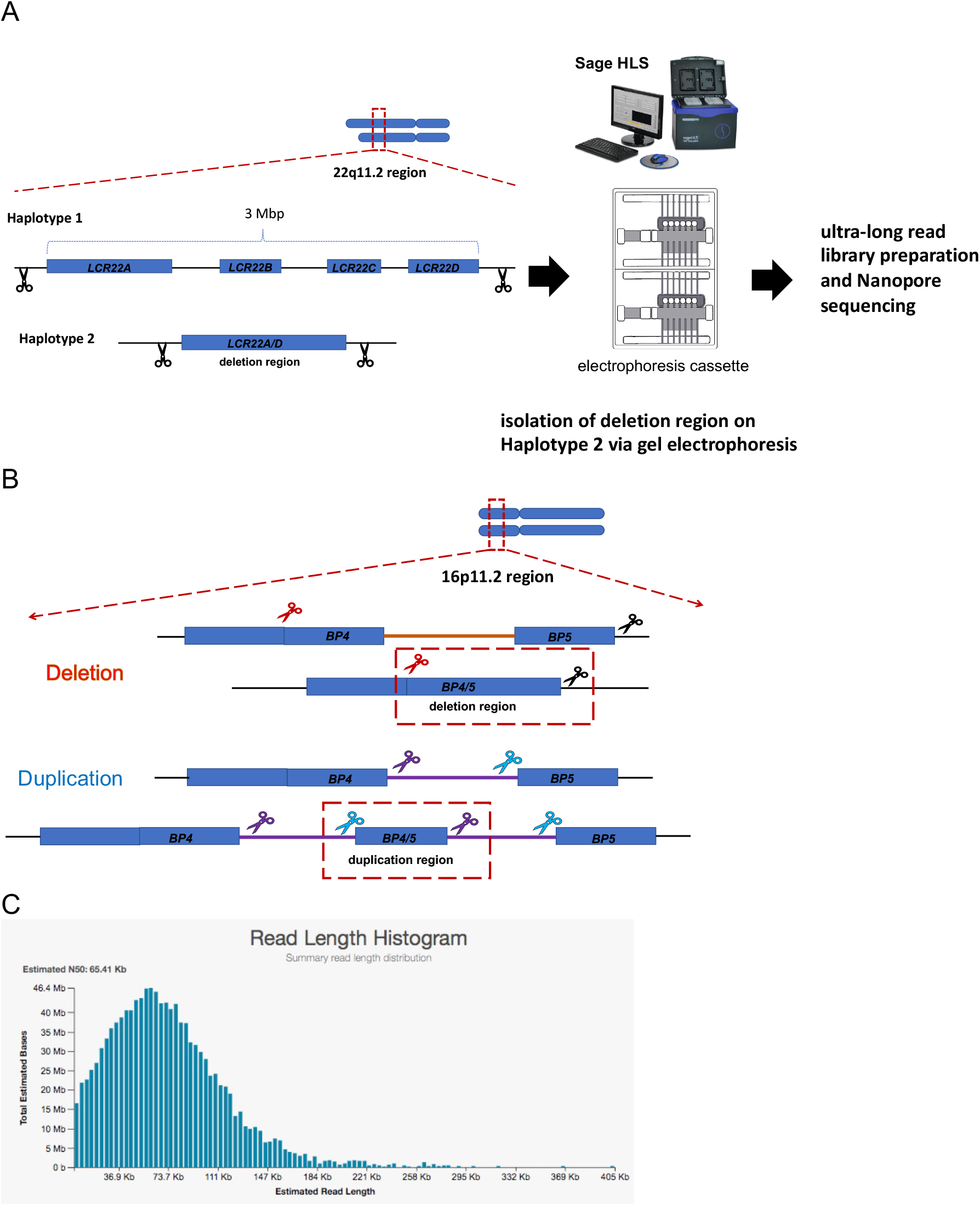
Method overview. (A) Experimental design of CTLR-Seq for targeting the 22q11.2 deletion region on Haplotype 2. The typical 22q11.2 deletion span ~3 Mbp spanning four major segmental duplication regions LCR22A-D. CRISPR gRNAs were designed to cut sites flanking the deletion region (Haplotype 2) that forms due to genomic rearrangement between LCR22A and LCR22D, concomitantly, sites that flank LCR22A and LCR22D respectively on Haplotype 1 are also cut. The 22q11.2 deletion region on Haplotype 2 is separated from Haplotype 1 and the rest of the genome via gel electrophoresis on the Sage HLS machine (Sage Science) followed by ultra-long read library preparation and Nanopore sequencing. (B) Genomic rearrangement of segmental duplication regions BP4 and BP5 give rise to 16p11.2 deletion and duplications. Design of CRISPR gRNA targets to isolate 16p11.2 BP4/5 deletion and duplication regions for CLR-Seq. Scissor colors represent different gRNA sequences. (C) Histogram of read Nanopore length distribution (22q11.2 deletion sample).

One of the key mysteries in psychiatric genetics is how such a wide range of clinical outcomes including non-penetrance can manifest from what is an ostensible issue of gene dosage. Attempts to model the biological effects of these CNVs have been bundled with challenges (Golzio and Katsanis, 2013). One of the main complicating factors is that the true extent of the genome rearrangements or “genomic scar” left by the CNVs, i.e. their *exact* boundaries or breakpoints, as well as their variability across different carriers have never been defined due to technological and methodological limitations. To truly understand the genetics and biological intricacies surrounding these CNVs, its essential to map out the details of the rearrangements at single base-pair resolution. In order to solve the sequence structure of the SegDup rearrangements that give rise to large recurring CNVs in the human genome, we applied CTLR-Seq to isolate intact, haplotype-specific target fragments (200-500 kb) of SegDup rearrangements. These target fragments are then used to generate targeted ultra-long Oxford nanopore sequencing libraries (Figure 1). The ultra-long sequencing reads are then used for *de novo* assembly.

## Materials and Methods

### Intact isolation of 200-500 kb genomic targets using CRISPR-TARGETED

Lymphoblastoid cells that harbor the CNV of interest were used as source of genomic DNA (gDNA). Approximately 375,000 live cells which yields approximately 2.5 micrograms of gDNA were used for each CRISPR-TARGETED experiment. Total DNA yield was measured using Qubit HS following the Mammalian White Blood Cell Suspension Kit (CELMWB1, Sage Science). CRISPR-TARGETED was performed on the Sage HLS machine (Sage Science) following manufacturer’s protocol (Manual-Rev-J-460034-12_12_19) using 100-300kb Target with 3 Hour Extraction and 4 Hour Separation. Target enrichment assay was performed using qPCR using RNase P as reference.

CRISPR gRNA pairs are the following at 6.5uM final concentration:

22q11.21DEL TGTAGCTACCTGTCGGCCTT GGGTCGTTTGGAATTGCGCT 16p11.2DEL GTTGATTTTCGTGCACGTGT TCTCCGGCTTGGTTCCTGCC 16p11.2DUP TGTTGATCTAAGTCGACCCG GTACTTTGAGATCACTTCCG

### Ultra-long read Nanopore library preparation

All steps were performed using wide-bore pipette tips. To purify DNA for library preparation, CRISPR-TARGETED elution containing >500,000 copies of targets (pooled from 2-4 experiments) were slowly transferred (<320 uL) to a 2mL Lo-Bind tube. DNA concentration must be <0.5ng/uL. Ampure XP (Beckman Coulter) beads (0.45X) were then added to the elution by ejecting beads from pipette tip placed at approximately 1 cm above the elution liquid surface. The mixture was mixed by gently tapping the bottom of the 2mL Lo-Bind tube (Eppendorf). The tube was then placed horizontally on a bench-top rotator and rotated at slowest speed for 15 min at room temperature. The tube was then placed on a magnetic rack and supernatant was removed once the beads were bound to the side of the tube; 2X ethanol wash (80%) was then performed. Beads were dried at room temperature for 1 minute and 50uL of EB buffer (Qiagen) was added slowly on top of the beads. The mixture was mixed by gentle tapping and placed on benchtop shaking heat block for 1 hour (37 degrees C, rotating at 400 rpm). The tube was then placed at 4 degrees C for 12 hours for elution. Ampure XP purification prior End-Repair and A-tail was not performed for the 22q11.21 deletion sample. Instead, elution was dialyzed on filter membrane for 1-2 hours to remove excess salt and then concentrated to 48 uL using SpeedVAC.

The elution mixture was placed on a magnetic rack, and once the solution is clear 48 uL of the elution was transferred to a 2mL Lo-Bind tube with End-Repair and A-tail buffer (7 uL) and enzymes (3 uL) premixed (NEBNext® Ultra™ II kit, New England Biolabs). The solution was then mixed by gentle tapping and transferred to a 0.2 mL PCR tube and incubated at 20 degrees C for 30 minutes and 65 degrees C for 30 minutes in a thermocycler; 25 uL of LNB buffer from LSK-109 kit (Oxford Nanopore Technologies) and 10 uL of ligase (New England Biolabs) and and 5 uL of adapter from LSK-109 kit was then added the reaction mixture and mixed by gentle tapping. The ligation reaction was then left at room temperature for 10 minutes. After ligation, the library was purified using the DNA purification procedure as described above except that the LFB buffer from the LSK-109 kit (Oxford Nanopore Technologies) was used to wash the XP beads (2 washes) where the tube was gently tapped to allow the XP beads to completely resuspend in the LFB buffer before placing on the magnet. After washing, the XP beads were dried for 30 seconds, and the final library was eluted with 12.5uL of EB buffer from the LSK-109 kit.

### Sequencing, alignment, and assembly

Libraries were sequenced for 48 hours on a MinION instrument (Oxford Nanopore Technologies) where library loading was conducted following standard manufacturer’s protocol. PASS filtered reads from MinKOWN were aligned to hg38 using minimap2 (parameters: -*ax map-pb*). On-target reads were identified as those that align to LCR22A and LCR22D regions for 22q11.2 deletion and BP4 and BP5 regions for 16p11.2 deletion and duplication. Manual overlap assembly of sequencing reads were performed using reads > 30 kb.

## RESULTS

### Intact isolation of 200-500kb target regions

The first part of the CTLR-Seq pipeline is the intact isolation of genomic targets interest. Targets can range from 50 kb to 1 Mb. Here, we demonstrate this application on three different targets of interest: the 22q11.2 deletion region and the 16p11.2 deletion and duplication regions (Figure 1). The typical 22q11.2 deletion span ~3 Mbp spanning four major segmental duplication regions, also known as low-copy repeats (LCR), LCR22A-D. CRISPR gRNAs were designed to cut immediately outside (inside unique genomic sequences) the segmental duplication rearrangements that form the deletion region (Figure 1A), concomitantly, sites that flank LCR22A and LCR22D respectively on haplotype without the genomic deletion are also cut. However, the excised fragment in this case will stretch approximately 3 Mbp, much larger than the expected size of target fragment containing the deletion region (few hundreds of kilobases). Due to this size difference, target fragment of interest containing the 22q11.2 deletion region separates from the other haplotype and the rest of the genome during gel-electrophoresis and isolated intact via electro-elution.

Similarly, CRISPR gRNAs were designed to cut immediately outside the 16p11.2 deletion and duplication regions. Although the 16p11.2 haplotype without the deletion or duplication (and other sites on the rearranged haplotype, in the case of the 16p11.2 duplication) will also be cut by the same gRNAs (Figure 1B), size differences between the excised fragments allow for the intact, haplotype-specific isolation of only the 16p11.2 deletion or duplication regions via gel-electrophoresis and electro-elution.

Target enrichment was assayed using qPCR where Taqman probes were designed to detect isolated targets of interest. Taqman probes for the other “normal” haplotype as well as also designed. We used a Taqman probe targeting the Rnase P gene to serve as reference. More than 200,000 targets of interest were obtained from a single assay for all three targets and the target enrichment relative to RNase P reference as greater than 30-fold on average. Background levels of the “normal” 22q11.2 and 16p11.2 haplotypes were the same as RNase P, indicating that the targets of interest were isolated in a haplotype-specific fashion. Approximating from where the target fragments of interest have migrated during gel-electrophoresis, sizes of the 22q11.2 deletion region, 16p11.2 deletion region, and 16p11.2 duplication regions were estimated to be 500 kb, 300 kb, and 200 kb respectively.

### Ultra-long read Nanopore sequencing

The second part of the CTLR-Seq pipeline is the preparation of ultra-long sequencing library (see Materials and Methods). Here, we show that even though the overall amount of DNA from a CRISPR-TARGETED elution is low, 16-40ng of DNA (0.2-0.5ng/uL), sequencing libraries up to 30-40x on-target coverage with N50 > 65 kb can still be achieved (Figure 1C, 2). For example, in the 22q11.2 deletion CTLR-Seq library, the N50 was 65.41 kb with the longest read spanning 405 kb; more than 80% of the reads span between 40 to 200 kb (Figure 1C). The 22q11.2 deletion CTLR-Seq library was prepared using an older version of the library preparation method where the elution was dialyzed first and then concentrated using speed vacuum. The on-target sequencing coverage obtained for the 22q11.2 deletion CTLR-Seq library is approximately 10x (Figure 2A). The 16p11.2 deletion and duplication CTLR-Seq libraries were prepared using the most updated CTLR-Seq library preparation method (see Materials and Methods) where approximately 30-40x on-target coverage was achieved (Figure 2B-C). The enrichment of target regions compared to background genomic regions is >40x for all CTLR-Seq libraries.

**Figure 2.**
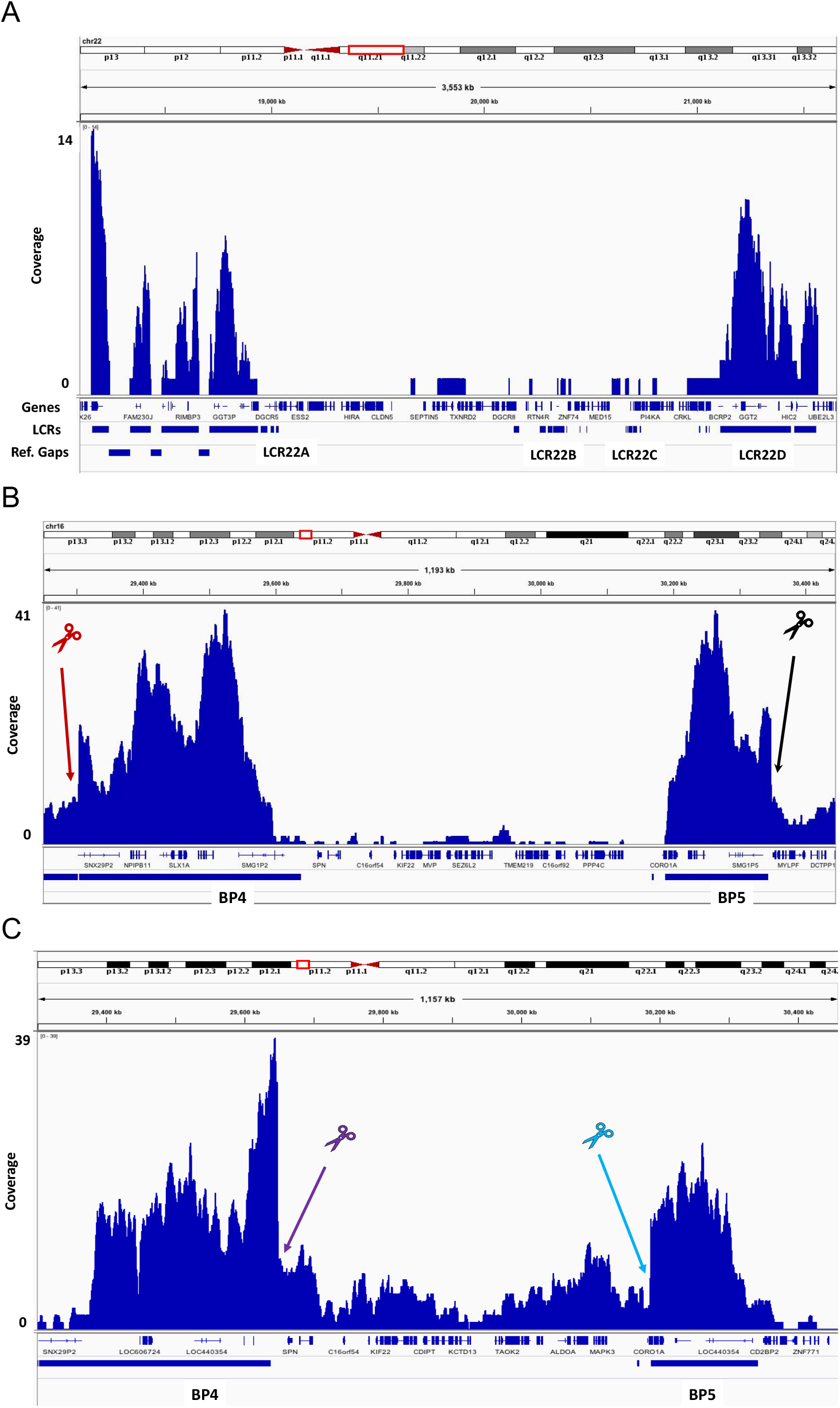
Sequencing coverage of 22q11.2 and 16p11.2 CNV regions for (A) 22q11.2 deletion, (B) 16p11.2 deletion, and (C) 16p11.2 duplication. Scissor colors represent different gRNA sequences.

### Assembly of segmental duplication rearrangement regions

The on-target reads obtained from CTLR-Seq typically span from 30 kb to 200 kb and are long enough to completely sequence assemble the target region of interest with manual overlapping assembly or other assembly tools built for handling Nanopore reads such as Flye (Kolmogorov et al., 2019), Canu (Koren et al., 2017), and wtdbg2 (Ruan and Li, 2020). Here, we performed manual overlap assembly by first identifying reads align to the 5’ and 3’ cutting sites of the gRNAs. We then use these reads as anchors to “stitch” together the assembly by finding overlapping reads. In the case of the 16p11.2 deletion and duplication regions, 2 to 3 long reads that span ~90 kb to ~200 kb is enough to assemble the entire target region of interest (Figure 3A-B). The 16p11.2 deletion assembly is 299.5 kb in length and can be completely assembled from two reads that span 102 kb and 226 kb respectively (Figure 3A). The 16p11.2 duplication assembly is 199.3 kb in length and can be completely assembled from 3 reads that all span ~90 kb (Figure 3B). The final sequence assembly size for both 16p11.2 deletion and duplication correspond well with approximately size of the enriched fragment from CRISPR-TARGETED. Both manual assemblies for the 16p11.2 CNV region were also obtained by using Flye (Kolmogorov et al., 2019) to assemble the on-target reads. The 22q11.2 deletion assembly was manually assembled by using 10 reads that span from 30 kb to 103 kb, and the total sequence assembly size for the 22q11.2 deletion is 484 kb (Figure 3C). This assembly was also orthogonally validated by using fiber-FISH optical mapping (Figure 3D). The probes used here for the fiber-FISH optical mapping analysis of the 22q11.2 region have been previously described (Demaerel et al., 2019).

**Figure 3.**
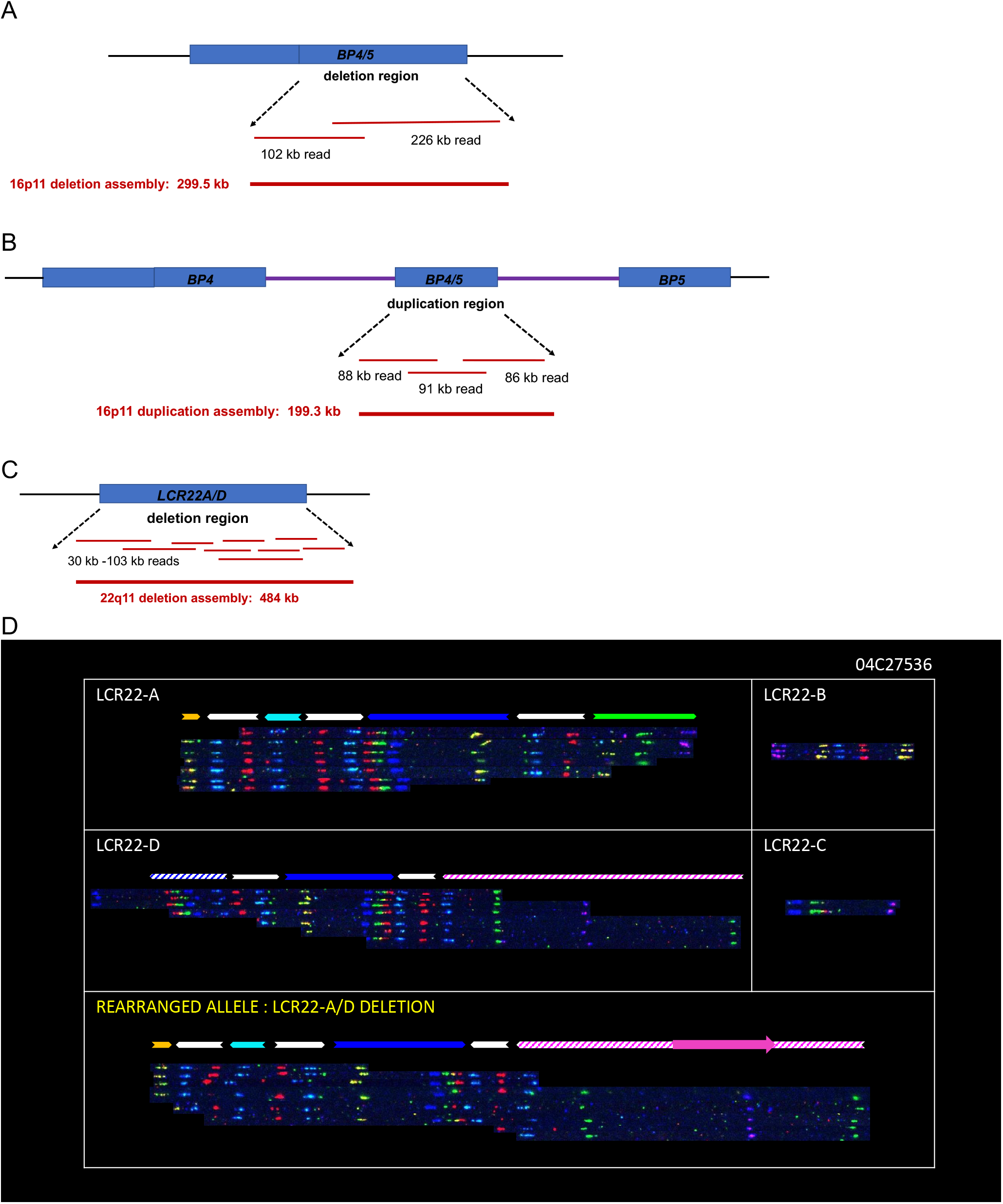
Sequencing assembly of 22q11.2 and 16p11.2 CNV regions for (A) 16p11.2 deletion, (B) 16p11.2 duplication, and (C) 22q11.2 deletion. The 16p11.2 deletion region (299.5 kb) is assembled from two reads (102 kb and 226 kb); the 16p11.2 duplication region (199.3 kb) is assembled from three reads (88 kb, 91 kb, and 86 kb), and the 22q11.2 deletion region is assembled from 9 reads ranging from 30 kb to 103 kb. (D) Fiber-FISH optical-based assembly for the 22q11.2 deletion haplotype (rearranged allele LCR22A/D deletion) and segmental duplication LCR22A-D for the other 22q11.2 haplotype without the genomic deletion.

## DISCUSSION

Here, we have developed a generally applicable method based on CRISPR/Cas9-targeted ultra-long read sequencing (CTLR-Seq) to completely and haplotype-specifically resolve, at base-pair resolution, large, complex, and highly repetitive genomic regions that had been previously impenetrable to next-generation sequencing analysis. To demonstrate its utility, we show, for the first time, the complete sequence assembly of the genome rearrangements that result in the 16p11.2 deletion and duplication CNVs and the 22q11.2 deletion. The important roles that these CNVs play in human developmental disorders has long been recognized (Marshall et al., 2008; Malhotra and Sebat, 2012; Kirov et al., 2014; Kirov, 2015; McDonald-McGinn et al., 2015; Niarchou et al., 2019); however, their sequences have never been resolved until now, despite decades of attempts (Morrow et al., 1995, 2018; Edelmann et al., 1999b; Shaikh et al., 2007; Guo et al., 2016). We also describe the development of a novel method CTLR-Seq which combines CRISPR-Cas9 and Nanopore long-read sequencing that allows for the targeted and haplotype-specific base-pair resolution of highly repetitive and complex segmental duplications that stretch hundreds of kilobases in the human genome (Bailey et al., 2002). Segmental duplications are known to be highly polymorphic across human populations for which the human reference genome is often contain gaps or misassembled (Bailey et al., 2001; Babcock et al., 2003; Nuttle et al., 2016; Demaerel et al., 2019). In such cases, *de novo* assembly using ultra-long reads as demonstrated here is preferable. This methodology also allows us to resolve the structures – equally unknown, residing in segmental duplications – of the other large pathogenic genomic rearrangements (i.e. on 1q21.1, 3q29, 15q11.2, 15q13.3, 16p11.2, and 16p13.1) (Malhotra and Sebat, 2012; Deshpande and Weiss, 2018; Takumi and Tamada, 2018). Other long and highly repetitive genomic sequences such as telomeres and centromeres may also be assembled and analyzed using CTLR-Seq.

The CTLR-Seq library preparation method takes advantage of the fact that the target genome fragments are isolated intact using CATCH (Jiang et al., 2015; Shin et al., 2017; Gabrieli et al., 2018). Since the genomic fragments are isolated intact and that the length Nanopore sequencing reads are limited by the size of input fragments, reductions in read lengths are typically the result of DNA shearing due to handling. Another key to generating successful ultra-long reads using CTLR-Seq is that the input DNA concentration must also be low (<0.5ng/uL). This low concentration makes it difficult to purify DNA using column or alcohol precipitation-based methods without suffering drastic yield loss, but allows for the use Ampure XP beads, from the library preparation reaction mixture and also from the CATCH elution buffer, which often contains high concentrations SDS that inhibits enzymatic activities during library prep. If the DNA concentration is high, then alcohol precipitation-based methods is then preferred since concentration of high molecular weight DNA will lead to the clumping of Ampure XP beads which drastically reduces purification yields. Currently, sequencing a single CTLR-Seq library on a Nanopore MinION flow-cell does not use up the entire sequencing capacity of the flow-cell. The utilization is only about 20-30%. In the future, pooling different barcoded libraries may be utilized to take full advantage of the sequencing capacity of Nanopore flow-cells.

A very important aspect of CTLR-Seq is that the use of CRISPR enables one to discriminate structural differences specific to a haplotype. This approach leverages the fact that some structural changes are large enough to enable isolation of different high molecular weight molecules representative of the separate haplotypes. Thus, more complex structure can be resolved efficiently. Because many complex structural variations are missed by even long-read whole genome sequencing (Shin et al., 2019), the greater coverage of complex genomic regions afforded by CTLR-Seq leads to the improved ability to resolve such complex structures. Finally, the targeted approach provided by CTLR-Seq to evaluating complex loci causative for specific diseases can be used to provide accurate resolution of these genomic structures within a diagnostic setting.

## Acknowledgements

We graciously acknowledge Dr. Chris Boles, CSO of Sage Science, for his technical support in operating the Sage HLS machine and associated workflows.

## References

Babcock M, Pavlicek A, Spiteri E, Kashork CD, Ioshikhes I, Shaffer LG, Jurka J, Morrow BE (2003) Shuffling of genes within low-copy repeats on 22q11 (LCR22) by Alu-mediated recombination events during evolution. Genome Res 13:2519–2532 Available at: http://www.ncbi.nlm.nih.gov/pubmed/14656960.

Bailey JA, Gu Z, Clark RA, Reinert K, Samonte R V, Schwartz S, Adams MD, Myers EW, Li PW, Eichler EE (2002) Recent segmental duplications in the human genome. Science 297:1003–1007 Available at: http://www.ncbi.nlm.nih.gov/pubmed/12169732.

Bailey JA, Yavor AM, Massa HF, Trask BJ, Eichler EE (2001) Segmental duplications: organization and impact within the current human genome project assembly. Genome Res 11:1005–1017 Available at: http://www.ncbi.nlm.nih.gov/pubmed/11381028.

Bernier R, Hudac CM, Chen Q, Zeng C, Wallace AS, Gerdts J, Earl R, Peterson J, Wolken A, Peters A, Hanson E, Goin-Kochel RP, Kanne S, Snyder LG, Chung WK, Simons VIP consortium (2017) Developmental trajectories for young children with 16p11.2 copy number variation. Am J Med Genet B Neuropsychiatr Genet 174:367–380 Available at: http://www.ncbi.nlm.nih.gov/pubmed/28349640.

Blumenthal I, Ragavendran A, Erdin S, Klei L, Sugathan A, Guide JR, Manavalan P, Zhou JQ, Wheeler VC, Levin JZ, Ernst C, Roeder K, Devlin B, Gusella JF, Talkowski ME (2014) Transcriptional consequences of 16p11.2 deletion and duplication in mouse cortex and multiplex autism families. Am J Hum Genet 94:870–883 Available at: http://www.ncbi.nlm.nih.gov/pubmed/24906019.

Demaerel W, Mostovoy Y, Yilmaz F, Vervoort L, Pastor S, Hestand MS, Swillen A, Vergaelen E, Geiger EA, Coughlin CR, Chow SK, McDonald-McGinn D, Morrow B, Kwok P-Y, Xiao M, Emanuel BS, Shaikh TH, Vermeesch JR (2019) The 22q11 low copy repeats are characterized by unprecedented size and structural variability. Genome Res 29:1389–1401 Available at: http://www.ncbi.nlm.nih.gov/pubmed/31481461.

Deshpande A, Weiss LA (2018) Recurrent reciprocal copy number variants: Roles and rules in neurodevelopmental disorders. Dev Neurobiol 78:519–530 Available at: https://pubmed.ncbi.nlm.nih.gov/29575775.

Deshpande A, Yadav S, Dao DQ, Wu Z-Y, Hokanson KC, Cahill MK, Wiita AP, Jan Y-N, Ullian EM, Weiss LA (2017) Cellular Phenotypes in Human iPSC-Derived Neurons from a Genetic Model of Autism Spectrum Disorder. Cell Rep 21:2678–2687 Available at: http://www.ncbi.nlm.nih.gov/pubmed/29212016.

Edelmann L, Pandita RK, Morrow BE (1999a) Low-copy repeats mediate the common 3-Mb deletion in patients with velo-cardio-facial syndrome. Am J Hum Genet 64:1076–1086 Available at: http://linkinghub.elsevier.com/retrieve/pii/S0002929707631735.

Edelmann L, Pandita RK, Spiteri E, Funke B, Goldberg R, Palanisamy N, Chaganti RS, Magenis E, Shprintzen RJ, Morrow BE (1999b) A common molecular basis for rearrangement disorders on chromosome 22q11. Hum Mol Genet 8:1157–1167 Available at: https://academic.oup.com/hmg/article-lookup/doi/10.1093/hmg/8.7.1157.

Gabrieli T, Sharim H, Fridman D, Arbib N, Michaeli Y, Ebenstein Y (2018) Selective nanopore sequencing of human BRCA1 by Cas9-assisted targeting of chromosome segments (CATCH). Nucleic Acids Res 46:e87–e87 Available at: https://academic.oup.com/nar/article/46/14/e87/4999242.

Ghebranious N, Giampietro PF, Wesbrook FP, Rezkalla SH (2007) A novel microdeletion at 16p11.2 harbors candidate genes for aortic valve development, seizure disorder, and mild mental retardation. Am J Med Genet A 143A:1462–1471 Available at: http://www.ncbi.nlm.nih.gov/pubmed/17568417.

Golzio C, Katsanis N (2013) Genetic architecture of reciprocal CNVs. Curr Opin Genet Dev 23:240–248 Available at: http://www.ncbi.nlm.nih.gov/pubmed/23747035.

Green Snyder L, D’Angelo D, Chen Q, Bernier R, Goin-Kochel RP, Wallace AS, Gerdts J, Kanne S, Berry L, Blaskey L, Kuschner E, Roberts T, Sherr E, Martin CL, Ledbetter DH, Spiro JE, Chung WK, Hanson E (2016) Autism Spectrum Disorder, Developmental and Psychiatric Features in 16p11.2 Duplication. J Autism Dev Disord 46:2734–2748 Available at: http://link.springer.com/10.1007/s10803-016-2807-4.

Guo X, Delio M, Haque N, Castellanos R, Hestand MS, Vermeesch JR, Morrow BE, Zheng D (2016) Variant discovery and breakpoint region prediction for studying the human 22q11.2 deletion using BAC clone and whole genome sequencing analysis. Hum Mol Genet 25:3754–3767 Available at: http://www.ncbi.nlm.nih.gov/pubmed/27436579.

Harel T, Lupski JR (2018) Genomic disorders 20 years on-mechanisms for clinical manifestations. Clin Genet 93:439–449 Available at: http://www.ncbi.nlm.nih.gov/pubmed/28950406.

Jiang W, Zhao X, Gabrieli T, Lou C, Ebenstein Y, Zhu TF (2015) Cas9-Assisted Targeting of CHromosome segments CATCH enables one-step targeted cloning of large gene clusters. Nat Commun 6:8101 Available at: http://www.nature.com/articles/ncomms9101.

Kaminsky EB et al. (2011) An evidence-based approach to establish the functional and clinical significance of copy number variants in intellectual and developmental disabilities. Genet Med 13:777–784 Available at: http://www.ncbi.nlm.nih.gov/pubmed/21844811.

Kirov G (2015) CNVs in neuropsychiatric disorders. Hum Mol Genet 24:R45–R49 Available at: https://pubmed.ncbi.nlm.nih.gov/26130694/.

Kirov G, Rees E, Walters JTR, Escott-Price V, Georgieva L, Richards AL, Chambert KD, Davies G, Legge SE, Moran JL, McCarroll SA, O’Donovan MC, Owen MJ (2014) The penetrance of copy number variations for schizophrenia and developmental delay. Biol Psychiatry 75:378–385 Available at: http://www.ncbi.nlm.nih.gov/pubmed/23992924.

Kolmogorov M, Yuan J, Lin Y, Pevzner PA (2019) Assembly of long, error-prone reads using repeat graphs. Nat Biotechnol 37:540–546 Available at: http://www.nature.com/articles/s41587-019-0072-8.

Koren S, Walenz BP, Berlin K, Miller JR, Bergman NH, Phillippy AM (2017) Canu: scalable and accurate long-read assembly via adaptive k -mer weighting and repeat separation. Genome Res 27:722–736 Available at: http://genome.cshlp.org/lookup/doi/10.1101/gr.215087.116.

Malhotra D, Sebat J (2012) CNVs: harbingers of a rare variant revolution in psychiatric genetics. Cell 148:1223–1241 Available at: http://www.ncbi.nlm.nih.gov/pubmed/22424231.

Marshall CR et al. (2008) Structural variation of chromosomes in autism spectrum disorder. Am J Hum Genet 82:477–488 Available at: http://www.ncbi.nlm.nih.gov/pubmed/18252227.

McCarthy SE et al. (2009) Microduplications of 16p11.2 are associated with schizophrenia. Nat Genet 41:1223–1227 Available at: http://www.ncbi.nlm.nih.gov/pubmed/19855392.

McDonald-McGinn DM, Sullivan KE, Marino B, Philip N, Swillen A, Vorstman JAS, Zackai EH, Emanuel BS, Vermeesch JR, Morrow BE, Scambler PJ, Bassett AS (2015) 22q11.2 deletion syndrome. Nat Rev Dis Prim 1:15071 Available at: http://www.ncbi.nlm.nih.gov/pubmed/27189754.

Migliavacca E et al. (2015) A Potential Contributory Role for Ciliary Dysfunction in the 16p11.2 600 kb BP4-BP5 Pathology. Am J Hum Genet 96:784–796 Available at: http://www.ncbi.nlm.nih.gov/pubmed/25937446.

Morrow B, Goldberg R, Carlson C, Das Gupta R, Sirotkin H, Collins J, Dunham I, O’Donnell H, Scambler P, Shprintzen R (1995) Molecular definition of the 22q11 deletions in velo-cardio-facial syndrome. Am J Hum Genet 56:1391–1403 Available at: http://www.ncbi.nlm.nih.gov/pubmed/7762562.

Morrow BE, McDonald-McGinn DM, Emanuel BS, Vermeesch JR, Scambler PJ (2018) Molecular genetics of 22q11.2 deletion syndrome. Am J Med Genet A 176:2070–2081 Available at: http://www.ncbi.nlm.nih.gov/pubmed/30380194.

Niarchou M, Chawner SJRA, Doherty JL, Maillard AM, Jacquemont S, Chung WK, Green-Snyder L, Bernier RA, Goin-Kochel RP, Hanson E, Linden DEJ, Linden SC, Raymond FL, Skuse D, Hall J, Owen MJ, Bree MBM van den (2019) Psychiatric disorders in children with 16p11.2 deletion and duplication. Transl Psychiatry 9:8 Available at: http://www.ncbi.nlm.nih.gov/pubmed/30664628.

Nuttle X et al. (2016) Emergence of a Homo sapiens-specific gene family and chromosome 16p11.2 CNV susceptibility. Nature 536:205–209 Available at: http://www.ncbi.nlm.nih.gov/pubmed/27487209.

Ruan J, Li H (2020) Fast and accurate long-read assembly with wtdbg2. Nat Methods 17:155–158 Available at: http://www.nature.com/articles/s41592-019-0669-3.

Sanders SJ et al. (2012) De novo mutations revealed by whole-exome sequencing are strongly associated with autism. Nature 485:237–241 Available at: http://www.ncbi.nlm.nih.gov/pubmed/22495306.

Sebat J et al. (2007) Strong association of de novo copy number mutations with autism. Science 316:445–449 Available at: http://www.ncbi.nlm.nih.gov/pubmed/17363630.

Shaikh TH, O’Connor RJ, Pierpont ME, McGrath J, Hacker AM, Nimmakayalu M, Geiger E, Emanuel BS, Saitta SC (2007) Low copy repeats mediate distal chromosome 22q11.2 deletions: sequence analysis predicts breakpoint mechanisms. Genome Res 17:482–491 Available at: http://www.ncbi.nlm.nih.gov/pubmed/17351135.

Shin G, Greer SU, Xia LC, Lee H, Zhou J, Boles TC, Ji HP (2019) Targeted short read sequencing and assembly of re-arrangements and candidate gene loci provide megabase diplotypes. Nucleic Acids Res 47:e115 Available at: http://www.ncbi.nlm.nih.gov/pubmed/31350896.

Shin G, Grimes SM, Lee H, Lau BT, Xia LC, Ji HP (2017) CRISPR–Cas9-targeted fragmentation and selective sequencing enable massively parallel microsatellite analysis. Nat Commun 8:14291 Available at: http://www.nature.com/doifinder/10.1038/ncomms14291.

Stankiewicz P, Lupski JR (2010) Structural variation in the human genome and its role in disease. Annu Rev Med 61:437–455 Available at: http://www.ncbi.nlm.nih.gov/pubmed/20059347.

Steinman KJ, Spence SJ, Ramocki MB, Proud MB, Kessler SK, Marco EJ, Green Snyder L, D’Angelo D, Chen Q, Chung WK, Sherr EH, Simons VIP Consortium (2016) 16p11.2 deletion and duplication: Characterizing neurologic phenotypes in a large clinically ascertained cohort. Am J Med Genet A 170:2943–2955 Available at: http://www.ncbi.nlm.nih.gov/pubmed/27410714.

Sullivan PF, Daly MJ, O’Donovan M (2012) Genetic architectures of psychiatric disorders: the emerging picture and its implications. Nat Rev Genet 13:537–551 Available at: http://www.ncbi.nlm.nih.gov/pubmed/22777127.

Takumi T, Tamada K (2018) CNV biology in neurodevelopmental disorders. Curr Opin Neurobiol 48:183–192 Available at: http://www.ncbi.nlm.nih.gov/pubmed/29331932.

Walsh KM, Bracken MB (2011) Copy number variation in the dosage-sensitive 16p11.2 interval accounts for only a small proportion of autism incidence: a systematic review and meta-analysis. Genet Med 13:377–384 Available at: http://www.ncbi.nlm.nih.gov/pubmed/21289514.

Ward TR, Zhang X, Leung LC, Zhou B, Muench K, Roth JG, Khechaduri A, Plastini MJ, Charlton C, Pattni R, Ho S, Ho M, Huang Y, Hallmayer JF, Mourrain P, Palmer TD, Urban AE (2020) Genome-wide molecular effects of the neuropsychiatric 16p11 CNVs in an iPSC-to-iN neuronal model. bioRxiv:2020.02.09.940965 Available at: http://biorxiv.org/content/early/2020/02/10/2020.02.09.940965.abstract.

Weiss LA et al. (2008) Association between microdeletion and microduplication at 16p11.2 and autism. N Engl J Med 358:667–675 Available at: http://www.ncbi.nlm.nih.gov/pubmed/18184952.

Zhang F, Gu W, Hurles ME, Lupski JR (2009) Copy number variation in human health, disease, and evolution. Annu Rev Genomics Hum Genet 10:451–481 Available at: http://www.ncbi.nlm.nih.gov/pubmed/19715442.

Zufferey F et al. (2012) A 600 kb deletion syndrome at 16p11.2 leads to energy imbalance and neuropsychiatric disorders. J Med Genet 49:660–668 Available at: http://www.ncbi.nlm.nih.gov/pubmed/23054248.

